# Filamin A mediates embryonical palatal fusion by linking mechanotransduction with β-Catenin/Smad2

**DOI:** 10.1101/2024.02.16.580664

**Authors:** Ziyi Wang, Satoru Hayano, Yao Weng, Xindi Mu, Mitsuaki Ono, Jeremie Oliver Piña, Rena N. D’Souza, Takashi Yamashiro, Toshitaka Oohashi, Hiroshi Kamioka

## Abstract

To decipher potential mechanisms underlying cleft palate (CP), we used advanced bioinformatic integrated with literature mining and genome-wide association study (GWAS). Re-analysis of RNA-seq data (GSE45568, GSE185279) combined with literature mining highlighted the roles of Filamin A (*Flna*) and Epithelial-Mesenchymal Transition (EMT) in palatal development. Immunofluorescence of *in vivo* palatal shelves showed increased *Flna* in medial edge epithelial (MEE) cells and EMT cells located in an epithelial triangle. Inhibition of TGF-β or RhoA and mechanical stimuli impacted *Flna* expression in *ex vivo* cultured palatal shelves. Re-analysis of scRNA-seq data (GSE155928) highlighted a correlation between *Flna* and *Ctnnb1* in EMT cells. *Flna* knockdown affected β-catenin/Smad2 expression in cultured palatal shelves and HaCaT cells. Epithelium-specific knockout of *Flna* delayed palatal fusion in female mice but not males. Mendelian randomization analysis suggested that parental habitual physical activity (HPA) was causally associated with lower risk of CP in their offspring. Together, these findings suggested that parental HPA could benefit their offspring’s palatal development through *Flna* by linking mechanotransduction with the Wnt/TGF-β/Smad signaling pathways during palatal fusion.

## Introduction

Understanding the mechanisms of maxillofacial development, particularly cleft palate, is crucial in orthodontics and dentistry for treating congenital disorders affecting the craniofacial complex. Specific to the maxilla, cleft palate, either isolated or combined with cleft lip, constitutes one of the most frequently diagnosed birth defects globally (Marazita, 2012). Interestingly, cleft lip alone and both cleft palate and lip are more common in males, and cleft palate alone is more common in females (Pool *et al*, 2021). While numerous hypotheses have been proposed to explain cleft palate, focusing on the adhesion and fusion of palatal shelves, the mechanisms of gender differences in cleft palate are poorly understand (Marazita, 2012; Lan *et al*, 2015; Nakajima *et al*, 2018; Tarr *et al*, 2018; Weng *et al*, 2018).

The palatal shelves mainly consist of mesenchymal cells derived from the cranial neural crest, covered by a single-layer embryonic epithelium, known as the medial edge epithelial (MEE) layer (Lan *et al*, 2015). Murine models have proven to be indispensable resources to study the developing in a mammalian system similar to that in humans. During normal palatal development, the epithelial fusion of palatal shelves in mice occurs around embryonic day 14 (E14), leading to the formation of a multilayered epithelial seam. This seam, created by the adhesion of opposing MEE cells at the palatal shelf tips, thins from E14.5 to E15.5 to a single-cell layer, allowing for mesenchymal integration of the palatal shelves to proceed (Piña *et al*, 2023). This process involves MEE cell migration-induced crowding and stretch receptor activation, initiating signaling pathways that result in epithelial cell extrusion, transforming the seam into distal epithelial triangles and medial epithelial islands (Nakajima *et al*, 2018; Tarr *et al*, 2018). Eventually, MEE cells are depleted, leaving only mesenchymal cells at the palatal midline (Nakajima *et al*, 2014). MEE cells are pivotal in palatal fusion, showing a preference for expressing transforming growth factor β (TGF-β) and receptor-regulated Smads (Nakajima *et al*, 2018; Lan *et al*, 2015). These Smads, including Smad1, Smad2, Smad3, Smad5, and Smad8/9, are directly regulated by TGF-β receptors. Additionally, Wnt/β-catenin signaling is essential for maintaining TGF-β expression in MEE cells during palatal fusion (Nakajima *et al*, 2018; Reynolds *et al*, 2020).

Cleft palate may manifest as part of Mendelian syndromes or as a clinical phenotype associated with chromosomal anomalies. For example, mutations in Filamin A (*FLNA*) lead to various severe malformations, including periventricular nodular heterotopia, otopalatodigital spectrum disorders, and sometimes cleft palate in humans (Fennell *et al*, 2015; Oshina *et al*, 2022). FLNA, the most abundant member of the filamin protein family, acts as a crosslinker for F-actin and interacts with many binding partners (Zhang *et al*, 2023). Its mechanosensing domain, responding to mechanical forces through conformational changes, is involved in various mechanotransduction signaling pathways, influencing cellular migration, morphology, and other crucial biological processes (Zhang *et al*, 2023, 21). One previous study has shown that FLNA-regulated apical epithelial extrusion squeezed out the oncogenic Src-transformed cells (Kajita *et al*, 2014). This type of epithelial cell extrusion, essential for removing damaged or undesirable cells, is also observed in epithelial triangles during palatal fusion (Nakajima *et al*, 2018). Recent studies indicate that FLNA regulates the Wnt/β-catenin and Smad signaling pathways in various types of cells (Yang *et al*, 2022; Cheng *et al*, 2020). However, the role of *FLNA* in embryonic palatal development remains unexplored.

Additionally, prenatal exposure to certain teratogens and environmental factors has been linked to an increased risk of cleft palate (Babai & Irving, 2023). For example, gestational exposure to air pollution can increase the relative risk of cleft palate by 10-28% (Huang *et al*, 2023). Moreover, women with physical disabilities are at a higher risk of delivering low birthweight and preterm infants (Morton *et al*, 2013), both known risk factors for cleft palate (Wyszynski *et al*, 2004).

Although it is known that multiple genetic and environmental etiologies contribute to cleft palate anomalies, few risk factors can be directly targeted in clinical practice. To address this, we used advanced bioinformatics combined with a systematic review of current literature on palate development and genome-wide association studies (GWAS) to identify a key mechanism of palatal fusion with the potential to reduce cleft palate risk through improved maternity care, a gene not previously prioritized. This study outlines a detailed bioinformatics-driven research workflow that illustrates how we gradually determined the role of *Flna* in regulating the Wnt/TGF-β/Smad signaling pathway during palatal fusion by following bioinformatics analysis clues. Specifically, the expression of X chromosome-linked *Flna* exhibited high sensitivity to even minor mechanical stimuli as a mechanosensor, linking mechanotransduction with the Wnt/TGF-β/Smad signaling pathways during palatal development. Supportive evidence of the role of mechanical stimuli in cleft palate came from large-scale GWAS, using Mendelian randomization (MR) analysis, which suggested a causality of parental habitual physical activity (HPA) in reducing cleft palate risk. We believe that this workflow can be widely applied in similar studies.

## Results

### *Flna* and Epithelial-Mesenchymal Transition are critical for palatal fusion

Two RNA-seq datasets were combined to identify overlapped differentially expressed genes (DEGs) and common themes in Gene Ontology Biological Process (GO_BP) terms across E12.5, E14.5, and E16.5 during palatal development. This revealed 39 DEGs, with a notable emphasis on “Migration” in GO_BP themes (**Fig. 1**, upper-right panel). Of these overlapped DEGs, five genes (*Mmp9*, *Anxa1*, *Sparc*, *Aqp3*, and *Adam12*) were significantly associated with the Medical Subject Headings (MeSH) term “Cell Movement” (**Fig. 1**, upper-right panel).

**Figure 1.**
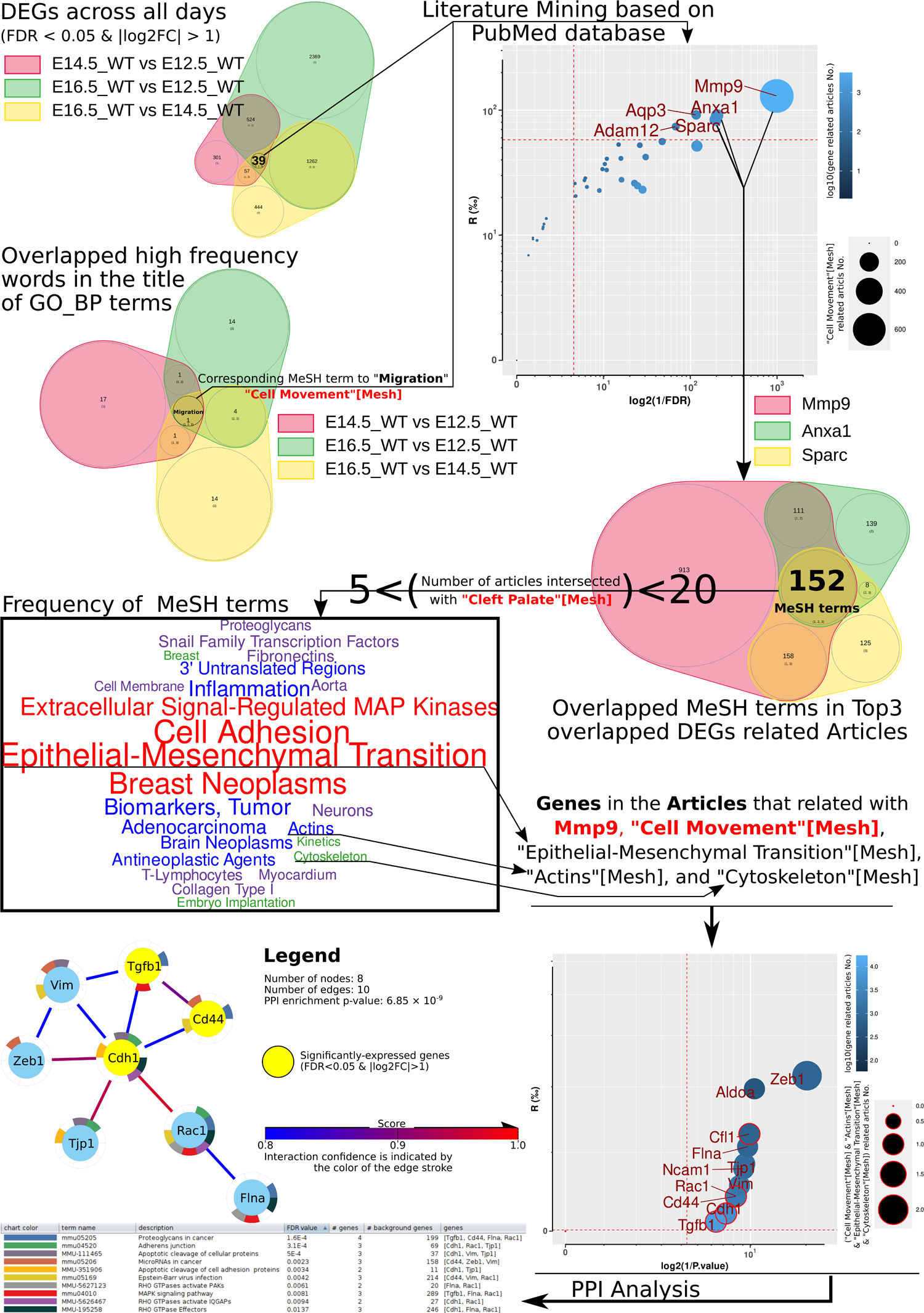
Re-analysis of previous bulk RNA-seq data combined with literature mining highlights Flna and Epithelial-Mesenchymal Transition in palatal fusion. By merging wild-type samples from two RNA-seq datasets, we identified 39 overlapped DEGs and a high-frequency word (“Migration”) in the GO_BP term. Literature mining revealed *Mmp9*, *Anxa1*, and *Sparc* as top genes associated with “Cell Movement”, corresponding to the MeSH term “Migration”. We analyzed 152 overlapped MeSH terms related to these genes, filtering out terms with either more than 20 (well-documented) or fewer than 5 (insufficient evidence) related articles, and calculated the frequency of remaining MeSH terms. Based on frequency and prior knowledge, we pinpointed “Epithelial-Mesenchymal Transition”, “Actins”, and “Cytoskeleton” as the top three frequent MeSH terms associated with *Mmp9* and “Cell Movement”. We then retrieved 11 genes linked to articles involving *Mmp9*, “Cell Movement”, “Epithelial-Mesenchymal Transition”, “Actins”, and “Cytoskeleton” from the PubMed Gene database. Among these, eight showed significant PPI. Notably, *Cdh1*, *Cd44*, and *Tgfb1* were also identified as DEGs in the combined dataset of GSE185279 and GSE109838, and these eight genes significantly enriched RHO GTPases/MAPK-related signaling pathways. **DEGs**, differentially expressed genes; **GO_BP**, Gene Ontology Biological Process; **GSE**, Genomic Spatial Event database; **MeSH**, Medical Subject Headings; **PPI**, Protein-Protein Interaction.

In detail, *Aqp3* and *Adam12* have limited reports regarding their association with human cleft palate. *Sparc* is involved in extracellular matrix and collagen turnover, with reduced levels observed in fibroblasts from cleft palate patients (Gagliano *et al*, 2010). *Anxa1* was identified as a key gene in mouse palatal fusion in another bulk RNA-seq dataset (GSE211601) and showed significant changes in an *in vitro* embryonic tissue fusion model using human cell lines (Cai *et al*, 2023; Belair *et al*, 2017). The single-nucleotide polymorphism (SNP) within *MMP9*, rs3918242, was considered a cleft palate risk factor, although GWAS did not support the role of rs3918242 in cleft palate (Letra *et al*, 2007; Kumari *et al*, 2019). However, increased MMP9-positive osteocytes in cleft palate patients’ alveolus/palate bone suggest a crucial downstream role of MMP9 in palatal development (Buile *et al*, 2022).

To identify key proteins linking *Mmp9*, *Anxa1*, *Sparc*, and cell movement, we analyzed overlapped MeSH terms in related articles. We filtered the 152 overlapping MeSH terms based on article counts, excluding terms with counts over 20 (well-documented) or under 5 (insufficient evidence). The remaining terms were ranked by frequency, as shown in **middle panel of Figure 1**. Given the complexity of palatal fusion tissue movement (Reynolds *et al*, 2020), we identified “Epithelial-Mesenchymal Transition”, “Actins”, and “Cytoskeleton” as the top MeSH terms related to collective cell migration (Rørth, 2009). We further filtered articles from these MeSH terms to ensure relevance to *Mmp9*, “Cell Movement” [Mesh], “Epithelial-Mesenchymal Transition” [Mesh], “Actin” [Mesh], and “Cytoskeleton” [Mesh]. This yielded a set of articles for further analysis. According to PubMed Gene Database annotations, we retrieved 11 genes discussed in these articles (**Fig. 1**, lower-right panel). Among these, eight genes showed significant protein-protein interactions (PPI), forming a PPI network (**Fig. 1**, lower-left panel). Except for *Tjp1*, all other genes have been linked to cleft palate. Notably, *Cdh1*, *Cd44*, and *Tgfb1* were also identified as DEGs in the combined dataset of GSE185279 and GSE109838.

These eight genes were significantly enriched in RHO GTPases/MAPK-related signaling pathways. MAPK signaling, a critical non-Smad pathway influenced by TGF-β signaling, plays a role in palatal fusion (Nakajima *et al*, 2018). Rho-Rac signaling, regulated by Flna, affects cell morphology and migration (Nakamura, 2013). Rac1-regulated fibronectin arrangement is essential in retinoic-acid-induced cleft palate (Tang *et al*, 2016). *Cdh1*, encoding E-Cadherin, is an epithelial marker and crucial in the Epithelial-Mesenchymal Transition (EMT) process (Tripathi *et al*, 2020). SNPs on *CD44* are associated with non-syndromic oral clefts (Park, 2005). *Vim*, a mesenchymal marker and EMT regulator (Tripathi *et al*, 2020), was identified as a cleft palate candidate gene in a GWAS study (Yan *et al*, 2020). Zeb1 inactivation rescues palatal fusion in Grhl2-deficient mouse embryos (Carpinelli *et al*, 2020). Additionally, Flna have also been identified as a cleft palate-associated gene in another RNA-seq dataset (GSE67525) (Potter & Potter, 2015).

### High expression of *Flna* in MEE cells and its correlation with cell morphology

*Flna* exhibited high expression in MEE cells from E14.5 to E15.5 (**Fig. 2A**). Notably, two CD90-positive cells within an epithelial triangle were observed, potentially indicating early-stage of EMT (**Fig. 2A**, dashed rectangle, and **Fig. 2B**). CD90 is a known marker for mesenchymal stem cells (Brown *et al*, 2019). Subsequent analysis revealed a higher *Flna* expression in epithelial cells compared to mesenchymal cells and, conversely, a higher CD90 expression in mesenchymal cells than in epithelial cells (**Figs. 2B1-3** and **Figs. 2C1-3**). Additionally, mesenchymal cells displayed lower circularity and solidity than epithelial cells (**Figs. 2B4-5** and **Figs. 2C4-5**). The two CD90-expressing cells, characterized by higher circularity and solidity, suggest an early stage of the EMT process in these cells (Tripathi *et al*, 2020). Moreover, there was a significant correlation between *Flna* and *CD90* expression, as well as a negative correlation of Flna expression with cell circularity and solidity in epithelial cells.

**Figure 2.**
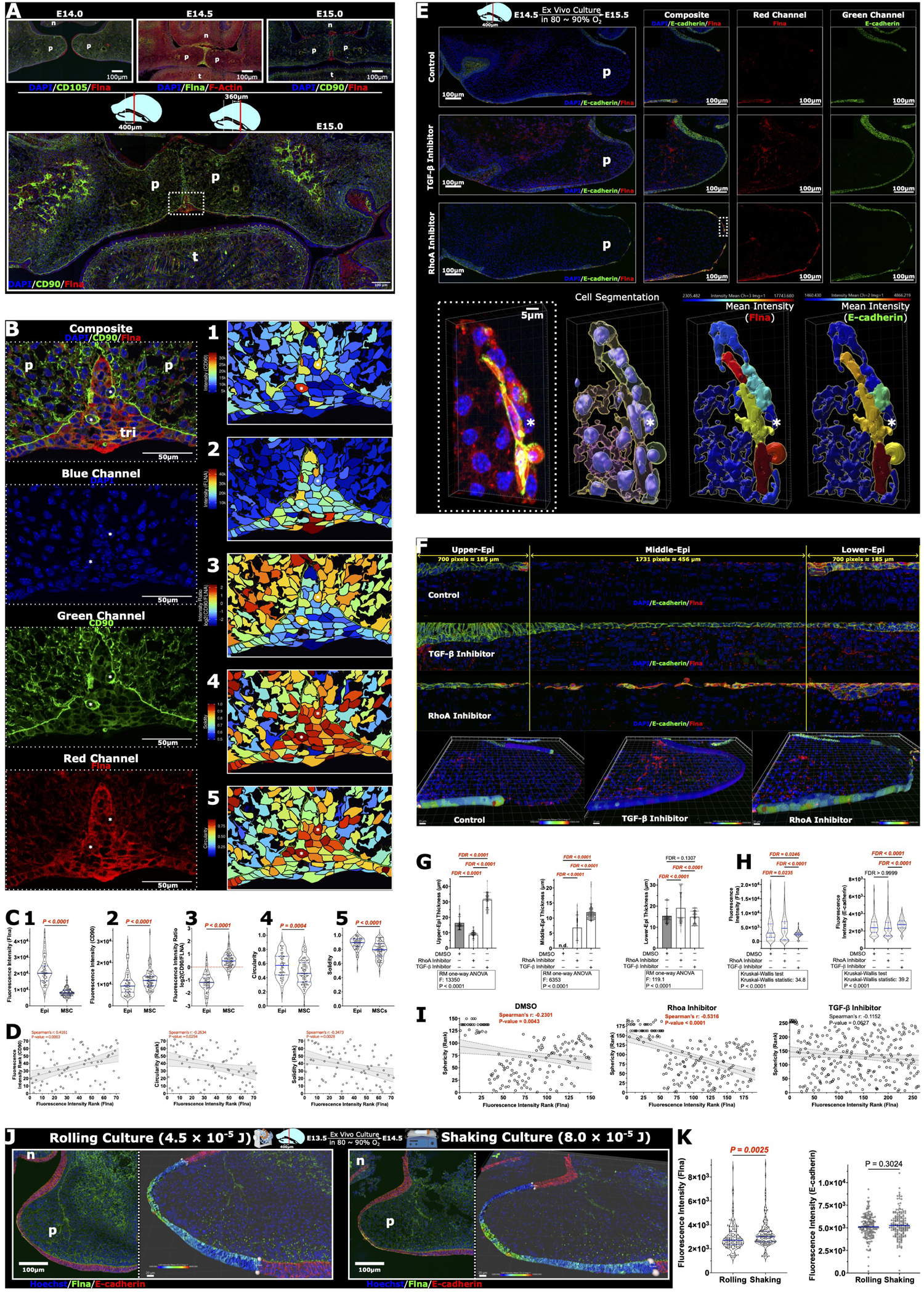
the correlation of Flna expression with MEE cells and its response to TGF-β and Rho signaling pathways, and mechanical stimuli. **A** In vivo sections of embryonic palatal shelves at E14, E14.5, and E15.0 showed a distinctive Flna expression pattern. At E15.0, two CD90-positive cells were observed within the epithelial triangle, highlighted in a dashed rectangle. **B** Fluorescence intensity quantification and cellular morphology analysis were conducted on these CD90-positive cells and their adjacent cells in the dashed rectangle area of (**A**) using an oil-immersed 63x objective lens. Pseudo-color images illustrate the spatial distribution pattern of Flna fluorescence intensity, CD90 fluorescence intensity, the fluorescence intensity ratio of CD90 to Flna, cellular circularity, and cellular solidity. **C** Quantitative measurements for Flna/CD90 fluorescence intensity and cell morphology correspond to images 1-5 in (**B**). **D** Spearman’s correlations were determined between Flna and CD90 expression, as well as cellular circularity/solidity in epithelial cells. E *Ex vivo* cultured palatal shelves treated with DMSO (0.1%), Y27632 (RhoA inhibitor; 50 μM), and SB431542 (TGF-β inhibitor; 20 μM) for 24 hours. The lower panel illustrates a Cdh1-negative cell surrounded by Cdh1-positive cells on the surface, induced by RhoA inhibitor treatment. **F** Straightening the epithelial layer from the images in (**E**) facilitates epithelial morphology quantification and cell segmentation. **G**, **H** Changes in epithelial morphology (**G**) and Flna/E-cadherin fluorescence intensity (**H**) in the palatal fusion region in response to RhoA/TGF-β inhibitors. **I** TGF-β inhibitor treatment disrupted the correlation between Flna expression and cell morphology. **J** Mechanical stimuli application to *ex vivo* cultured palatal shelves and the analysis method. The right panel displays the palatal MEE cells region of interest and the resulting cell segmentation. **K** Quantification of Flna/E-cadherin fluorescence intensity for each cell was performed using the image from (**J**). **Data information:** For better presentation, all fluorescence images were uniformly adjusted for contrast, gamma, and brightness. All 2D images are after 3D projection. Quantitative results are shown in scatter or violin plots with individual data points, mean ± standard deviation for normally distributed data, or median with interquartile range for non-normally distributed data. All regression lines include 95% confidence intervals. Scale bars: 100 μm in (**A**), upper panel of (**E**), and (**J**); 20 μm in (**F**); 50 μm in (**B**). **p**, palatal shelves; **n**, nasal septum; **t**, tongue; **tri**, epithelial triangle; **RM one-way ANOVA**, repeated measures one-way analysis of variance.

**Figure 3.**
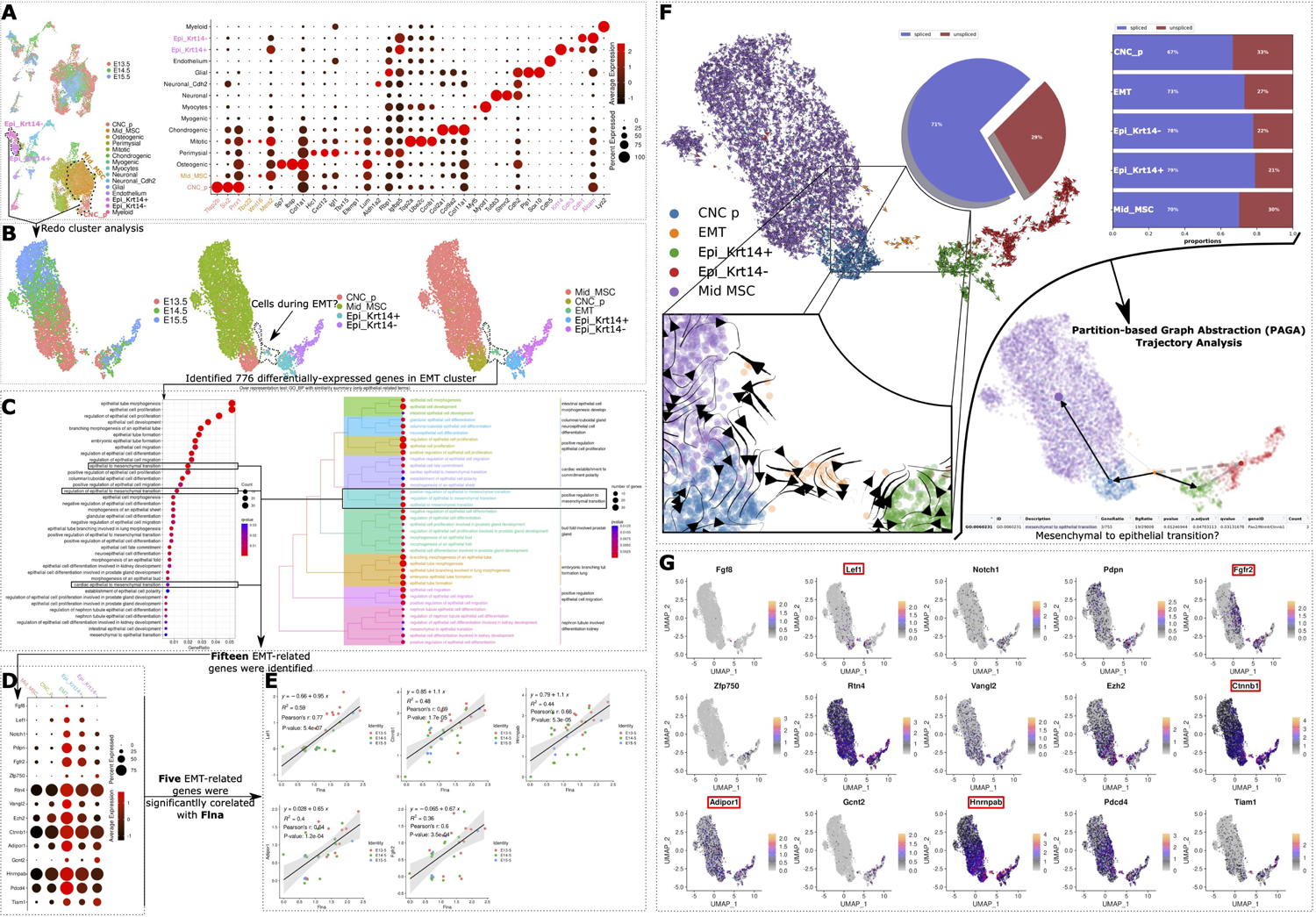
Re-analysis of single-cell RNA-seq dataset revealed EMT cell cluster and correlations of EMT markers with Flna expression during palatal development. **A** Wild-type samples from GSE155928 were re-analyzed, and cell type annotations were redone using markers reported in the original article. **B** By re-clustering subsets of CNC_P, Mid_MSC, Epithelium, and Epithelium_Cdh1, we increased clustering resolution and identified an EMT cell cluster. **C** Cell markers in the EMT cluster, identified as **DEGs** compared to other clusters, were enriched in EMT-related GO_BP terms. **D** These enriched EMT-related GO_BP terms included 15 cell markers that were highly expressed in the EMT cluster relative to Mid_MSC, CNC_P, Epi_Krt14+, and Epi_Krt14-clusters. **E** Among these 15 EMT-related markers, the expression of 5 markers was correlated with Flna expression in the EMT cluster. **F** Trajectory analysis based on RNA velocity were performed. **G** Expression patterns of all 15 EMT-related cell markers are shown, with Flna-related EMT markers highlighted in red rectangles. **Data information:** All regression lines are presented with a 95% confidence interval. **CNC_p,** cranial neural crest (CNC)-derived mesenchymal progenitors; **Mid_MSC**, midline mesenchymal cells; **EMT**, epithelial-mesenchymal transition; **Osteogenic**, osteogenic cells; **Perimysial**, perimysial cells; **Mitotic**, mitotic cells; **Neuronal**, neuronal cells; **Neuronal_Cdh2**, Cdh2-positive neuronal cells; **Glial**, glial cells; **Endothelium**, endothelial cells; **Epi_Krt14+**, Krt14-positive epithelial cells; **Epi_Krt14-**, Krt14-negative epithelial cells; **GO_BP**, the biological process of Gene Ontology term; **DEGs**, differential expressed genes.

### Impact of TGF-β and RhoA inhibitors on palatal fusion and association with *Flna* expression

Both the TGF-β and RhoA inhibitors dramatically disrupted the disappearance of the MEE during palatal fusion compared to the dimethyl sulfoxide (DMSO) control group (**Fig. 2e**). Interestingly, an E-cahderin-negative cell was observed surrounded by highly E-cahderin-enriched cells (highlighted with an asterisk in the bottom panel of **Fig. 2e**). This cell is either a mesenchymal cell that migrated into the remnant epithelium layer or a cell in the process of EMT. Given the critical role of RhoA in regulating cell migration(Nakamura, 2013) and EMT(Miyakawa *et al*, 2022), these processes were likely disrupted in the palatal shelves treated with the RhoA inhibitor. Thus, the observed cell (asterisk in **Fig. 2e bottom panel**) is suspected to be arrested during the EMT process.

Furthermore, as shown in **Figure 2F**, we measured epithelial morphology after straightening the epithelial layer of the palatal shelves. In the control group, where epithelial disappearance was observed, the remaining epithelium was segmented into three areas: upper remaining epithelium (Upper-Epi), middle remaining epithelium (Middle-Epi), and lower remaining epithelium (Lower-Epi). The thickness of the epithelium was measured pixel-by-pixel horizontally along the straightened layer. Our findings demonstrated that the RhoA inhibitor significantly reduced the mean thickness of the Upper-Epi but increased it in the Middle- and Lower-Epi when compared to the control group (**Fig. 2G**). On the other hand, the TGF-β inhibitor notably augmented the mean thickness of the Upper- and Middle-Epi, with no significant change in the Lower-Epi relative to the control group (**Fig. 2G**). Using a semi-automatic segmentation algorithm from Bitplane Imaris software, we identified epithelial cells in the Upper- and Lower-Epi (**Fig. 2F** bottom panel). The mean fluorescence intensity of Flna in each cell was significantly increased with the RhoA inhibitor and decreased with the TGF-β inhibitor, compared to the control group (**Fig. 2H** left panel). Meanwhile, the mean fluorescence intensity of E-cadherin in each cell was significantly increased by the TGF-β inhibitor, with no significant changes following RhoA inhibitor treatment compared to the control group (**Fig. 2H** right panel).

Significant negative Spearman’s correlations between *Flna* expression and cellular morphology were observed in the DMSO control and RhoA inhibitor groups but not in the TGF-β group (**Fig. 2I**). Because Flna has been reported to regulate cell morphology directly (Nakamura, 2013), our results indicated that Flna-regulated cellular morphological changes were dependent on TGF-β signaling.

### Mechanical stimuli increased *Flna* expression in MEE cells

As shown in **Fig. 2G**, the mean epithelium thickness of the Upper- and Lower-Epi in the TGF-β inhibitor group is approximately 23 µm. In the mechanical stimuli experiment on the *ex vivo* cultured embryonic palatal shelves, the Middle-Epi was defined as the thickness of epithelium less than 25 µm (**Fig. 2J**). The cells within the Middle-Epi were identified using Bitplane Imaris software, as described in the previous section. Interestingly, the mechanical stimuli significantly increased the expression of *Flna* in the early stage of palatal development, while there was no significant effect on the expression of E-cadherin (**Fig. 2K**).

### Re-analysis of single-cell RNA-seq dataset revealed *Flna*-EMT axis during palatal development

We performed clustering-based cell annotation using the cell markers reported by the original study (**Fig. 3A**) (Han *et al*, 2021). To achieve a higher clustering resolution, a subset containing only mesenchymal cells (cranial neural crest-derived mesenchymal cells, CNC_p; midline mesenchymal cells, Mid_MSC) and epithelial cells (Krt14-positive, Epi_Krt14+; Krt14-negative, Epi_Krt14-) was selected for detailed analysis. A distinct cluster (demarcated by a dashed line in **Fig. 3B**), encompassing all four cell types (CNC_P, Mid_MSC, Epi_Krt14+, Epi_Krt14-), emerged between the mesenchymal and epithelial.

Within this cluster, 776 DEGs were identified compared to all other clusters. These DEGs were notably enriched in GO_BP terms associated with EMT, specifically in terms of “epithelial to mesenchymal transition” (GO:0001837), “regulation of epithelial to mesenchymal transition” (GO:0010717), and “positive regulation of epithelial to mesenchymal transition” (GO:0010718), encompassing 15 of the 776 DEGs (**Fig. 3C**). These 15 DEGs, identified as EMT markers, were highly expressed in the EMT cell cluster (**Fig. 3D**).

Among these EMT markers, seven have critical roles in cleft palate pathogenesis. *Fgf8* and *Fgfr2* are involved in FGF signaling; *Lef1* and *Ctnnb1* are involved in Wnt signaling; both pathways are well-documented in palatal development and cleft palate (Tripathi *et al*, 2020; Nakajima *et al*, 2018). *EZH2* mutations relate to Weaver Syndrome (Gibson *et al*, 2012), often presenting with cleft palate (Tatton-Brown *et al*, 2013). *Notch1* may contribute to all-trans retinoic acid (ATRA)-induced cleft palate in mice (Zhang *et al*, 2016), and HNRNPAB is linked to Ankyloblepharon-Ectodermal Defects-Cleft Lip and/or Palate Syndrome by regulating FGFR2 (Fete *et al*, 2009). Notably, *Ctnnb1*, *Lef1*, *Fgfr2*, and *Hnrnpad* exhibited significant correlations with *Flna* (**Fig. 3E**).

RNA velocity-based trajectory analysis suggested potential differentiation paths from the EMT cluster to CNC_P and epithelial cells (**Fig. 3F**). This aligns with the enrichment of the DEGs in the “mesenchymal to epithelial transition” GO_BP term (**Fig. 3C**), suggesting mesenchymal-to-epithelial transition (MET) during palatal development. Although previous studies have demonstrated that glucocorticoids can induce both cleft palate and MET (Andrew & Zimmerman, 1971; Zhang *et al*, 2010), the role of MET in palatal development remains understudied.

The distribution of these 15 EMT markers was confirmed using Uniform Manifold Approximation and Projection (UMAP) plots (**Fig. 3G**). The UMAP plot analysis (**Figs. 3B** and **G**) showed that mesenchymal cells (CNC_p and Mid_MSC) and epithelial cells (Epi_Krt14+ and Epi_Krt14-) could be differentiated based on UMAP_1 values, while UMAP_2 values distinguished between stages of cell differentiation. The putative EMT cell cluster, characterized by intermediate UMAP_1 and lower UMAP_2 values, indicates their transitional state between mesenchymal and epithelial cells. This supports the hypothesis that these cells are in an EMT state, further validated by the significant enrichment of EMT-related GO_BP terms among their DEGs.

### Effects of *Flna* knockdown on epithelial disappearance, β-Catenin/Smad2 expression, and cell morphology in MEE cells of *ex vivo* cultured palatal shelves

In *ex vivo* cultured palatal shelves, siRNA-mediated *Flna* knockdown disrupted palatal fusion by impeding the disappearance of MEE cells, observable in both 48-hour and 24-hour cultures (**Figs. 4A** and **B**). Given that the lower remaining epithelium (Lower-Epi) typically disappears at a later stage, as shown in **Figure 4A** and supplementary images (deposited at Figshare, see **Reagents and Tools Table**), we focused on the upper remaining epithelium (Upper-Epi) to assess the efficacy of siFlna transfection. As demonstrated in the control group of **Figure 3E**, cells with high *Flna* expression were localized to a small area at the end of the Upper-Epi. Therefore, we selected epithelial cells within a 77 µm range from the end of the Upper-Epi for detailed analysis of epithelial and MEE cell morphology (**Fig. 4C**). To ensure consistent fluorescence intensity across different biological replicates and batches, we applied Min-Max normalization.

**Figure 4.**
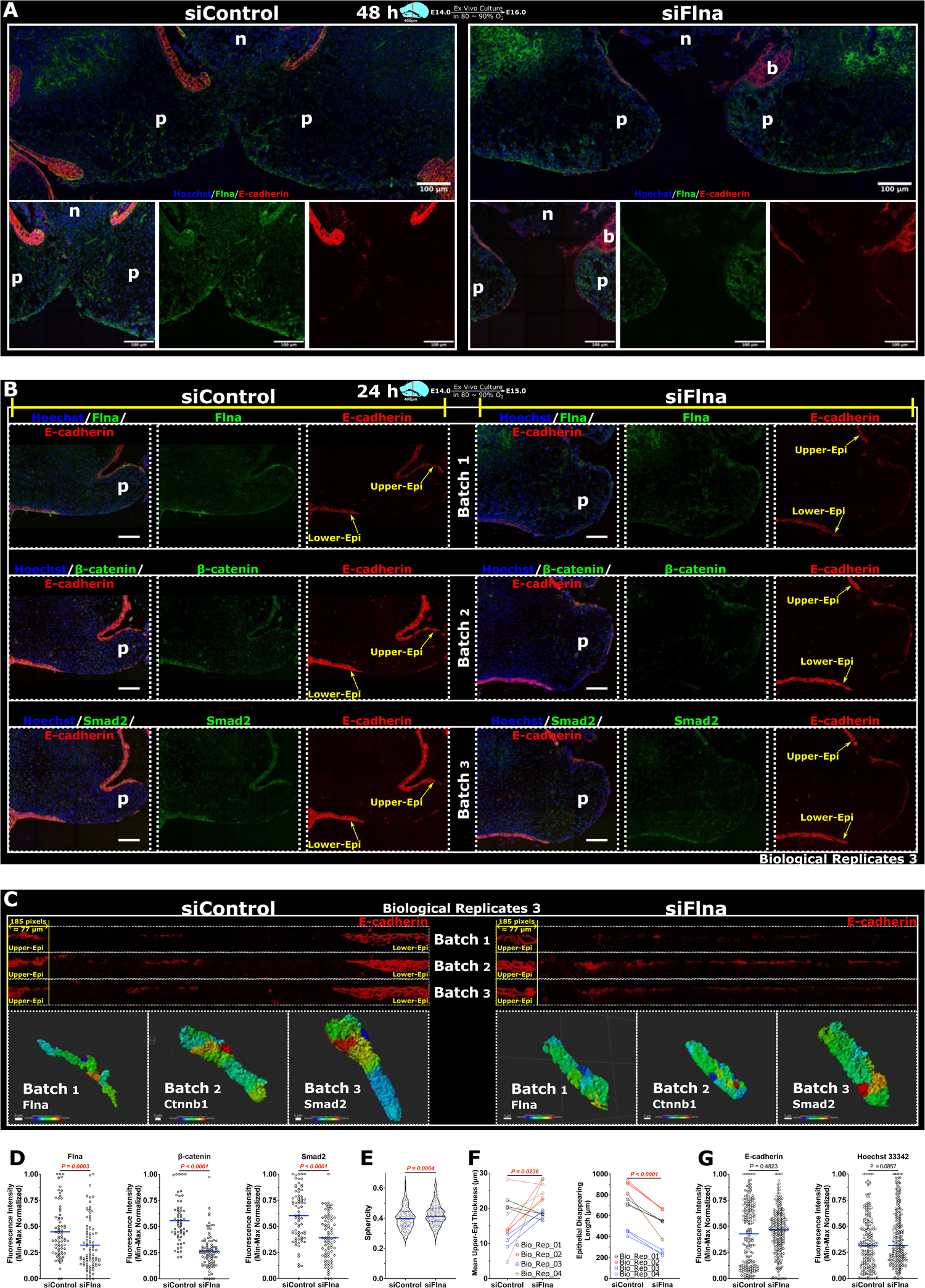
Effects of Flna knockdown on epithelial disappearance, β-Catenin/Smad2 expression, and cell morphology in MEE cells of ex vivo cultured palatal shelves. **A** *Ex vivo* cultured palatal shelves after transfection of siFlna using electroporation for 48 hours. Images on the upper panel were from a 40x objective lens, and images with separated fluorescence channels were from an oil-immersed 63x objective lens. **B** Representative results of Flna knock-down and experiment design for biological replicates. There are four biological replicates from 4 different pregnant mice in this study. **C** Straightening the epithelial layer from images of (**B**) shows the method for quantification of epithelial morphology and the epithelial part for cell segmentation in the upper epithelial region. **D, E, F** Changes in (**D**) fluorescence intensity of Flna, β-catenin, and Smad2; (**E**) cellular sphericity; (**F**) the amount of remaining epithelium after electroporation of siFlna on four biologically replicated *ex vivo* cultured embryonic palatal shelves. **Data information:** The contrast, gamma, and brightness of all fluorescence images were uniformly adjusted for better presentation. All 2D images here are images after 3D projection. Quantitative results are presented by scatter or violin plots showing individual data points along with mean ± standard deviation for data with normal distribution or median with interquartile range for data without normality. The linked lines of paired data in (**F**) were presented for the paired t-test. **p**, palatal shelves; **n**, nasal septum; **b**, burning damage due to the electroporation; scale bar = 100 μm. The contrast, gamma, and brightness of all fluorescence images were uniformly adjusted for better presentation.

Quantitative analysis of immunofluorescence images from embryos of four different pregnant mice showed a significant decrease in the protein level of Flna, β-catenin, and Smad2 in the Upper-Epi cells of the siFlna groups (**Fig. 4D**). Consistent with findings from *in vivo* samples (**Figs. 2B-D**) and *ex vivo* samples treated with inhibitors (**Figs. 2E-I**), *Flna* knockdown significantly increased cellular sphericity in Upper-Epi cells (**Fig. 4E**). Additionally, *Flna* knockdown increased the mean epithelial thickness in the Upper-Epi and the length of remaining epithelium in the Middle-Epi (**Fig. 4F**). These morphological alterations induced by *Flna* knockdown showed a similar tendency of the changes observed in TGF-β inhibitor-treated palatal shelves (**Figs. 2E-H**), suggesting that Flna influences palatal fusion *via* the Wnt/Smad-dependent TGF-β signaling pathway. The median fluorescence intensity of E-cadherin and Hoechst remained unchanged between siControl and siFlna groups, indicating that the Min-Max normalization effectively neutralized technical variations (**Fig. 4G**).

### Knockdown of *Flna* in HaCaT cells affects the expression and phosphorylation of β-catenin/SMAD2 and their response to TGF-β signaling

In HaCaT (human keratinocyte) cells, treatment with TGF-β3 significantly upregulated Flna protein levels, a response independent of Rho signaling, as evidenced by the lack of additional impact from RhoA inhibitor (**Figs. 5A** and **B**). This pattern aligns with previous observations in embryonic palatal MEE cells (**Figs. 2E-H**, and **5B**). *FLNA* knockdown by siRNA effectively reduced FLNA protein levels under various experimental conditions in HaCaT cells (**Fig. 5B**).

**Figure 5.**
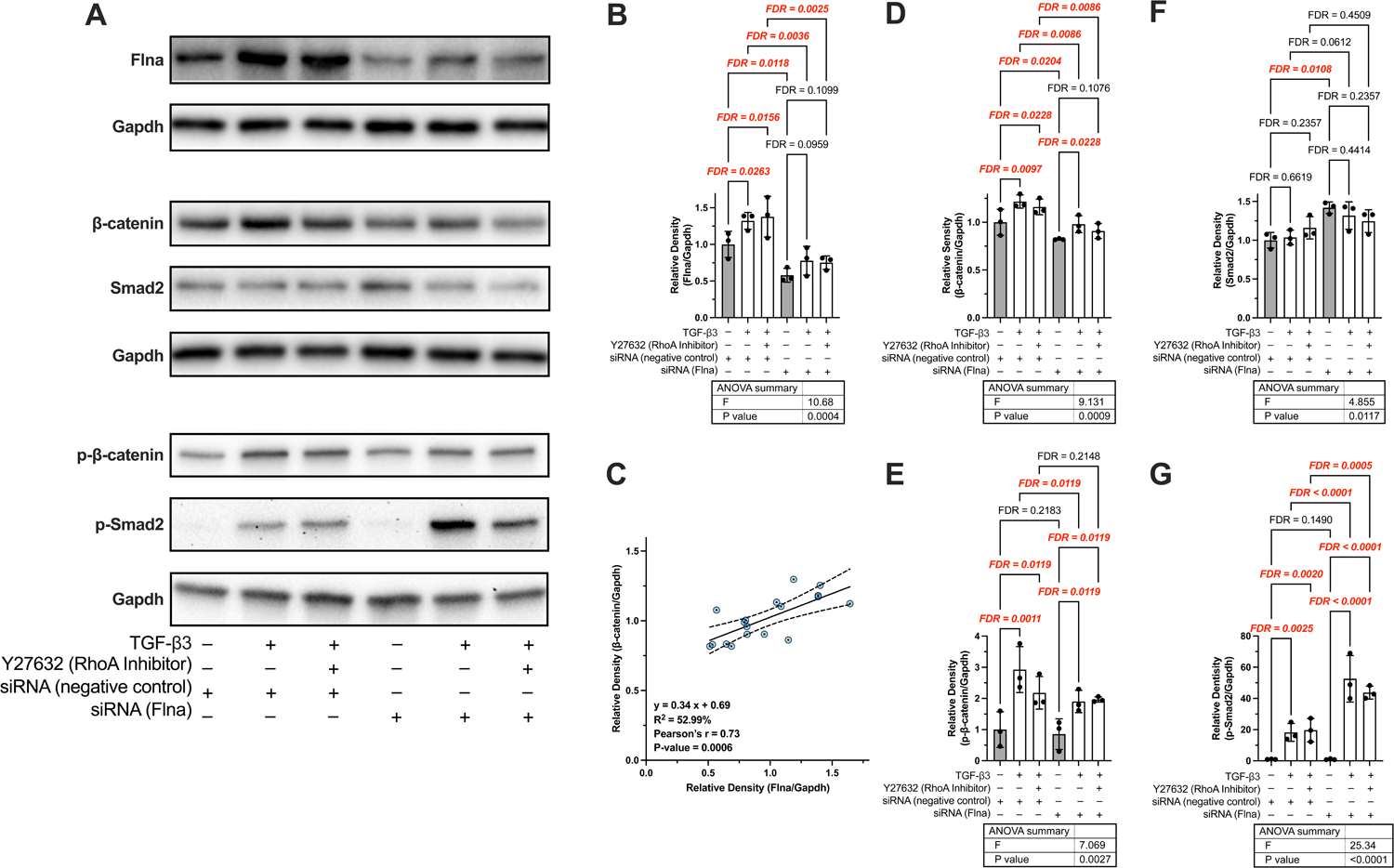
Knockdown of Flna in the HaCaT cell affected the expression and phosphorylation of β-catenin/Smad2 and their response to TGF-β signaling. **A** Representative results of Western Blotting. **B, D, E, F** The relative density of (**B**) FLNA, (**D**) β-catenin, (**E**) p-β-catenin, (**F**) SMAD2, and (**G**) p-SMAD2 was quantified using the Western Blotting results from three biological replicates. **C** Correlations between the relative density of FLNA and β-catenin. Quantitative results are presented by scatter plots showing individual data points along with mean ± standard deviation. All regression lines are presented with 95% confidence interval. **Data information: p-β-catenin**, phosphorylated β-catenin on ser552; **p-Smad2**, phosphorylated SMAD2 on Ser465/467. ANOVA, one-way analysis of variance.

Analyses shown in **Figures 5C** and **3E** reveal a significant correlation between FLNA and β-catenin in HaCaT cells and EMT cells *in vivo*. The strength of this correlation was remarkably similar, both at the protein level in HaCaT cells (Pearson’s *r*: 0.73; R^2^: 0.52) and gene level in scRNA-seq data (Pearson’s *r*: 0.69; R^2^: 0.48).

TGF-β3 treatment increased β-catenin levels in both siControl and siFlna HaCaT cells (**Fig. 5D**). However, the RhoA inhibitor disrupted this TGF-β3-induced β-catenin increase in siFLNA but not in siControl cells, suggesting necessity of RhoA for TGF-β3-enhanced β-catenin in the absence of FLNA (**Fig. 5D**). Additionally, TGF-β3 treatment, with or without the RhoA inhibitor, increased β-catenin phosphorylation at Ser552 in both siControl and siFLNA cells (**Fig. 5E**). While FLNA knockdown did not alter β-catenin phosphorylation, it significantly reduced TGF-β3-induced β-catenin phosphorylation to levels comparable to those observed with combined TGF-β3 and RhoA inhibitor treatment in siFLNA cells (**Fig. 5E**). This suggests that influence of RhoA on TGF-β3-induced β-catenin phosphorylation at Ser552 is FLNA-dependent in HaCaT cells.

Interestingly, only *FLNA* knockdown increased Smad2 protein levels, with no effects observed from either TGF-β3 or RhoA inhibitor treatments (**Fig. 5F**). However, phosphorylation of SMAD2 at Ser465/467 was enhanced by both TGF-β3 alone and in combination with the RhoA inhibitor in both siControl and siFLNA cells (**Fig. 5G**). *FLNA* knockdown did not affect SMAD2 phosphorylation per se, but it significantly amplified TGF-β3-induced phosphorylation of SMAD2 (**Fig. 5G**). These findings indicate that the effect of FLNA on TGF-β3-induced SMAD2 phosphorylation at Ser465/467 in HaCaT cells is independent of RhoA.

### Epithelium-specific knockout of *Flna* delayed palatal fusion in female mice but not males

We obtained 6 control females (*Flna^fl/f^*^l^), 4 *KRT14-Cre/+;Flna^fl/fl^*, 4 control male (*Flna^fl/y^*), and 9 *KRT14-Cre/+;Flna^fl/y^* E14.5 embryos from two pregnant mice, and 4 *Flna^fl/fl^*, 5 *KRT14-Cre/+;Flna^fl/fl^*, 4 *Flna^fl/y^*, and 4 *KRT14-Cre/+;Flna^fl/y^* E15.5 embryos from three pregnant mice (**Genotyping_Results.pdf** in **Mendeley Data**, see **Reagents and Tools Table**). **Figure 6A** shows representative results of Flna-cKO in female/male anterior/middle palatal shelves at E14.5 and E15.5; the analysis method is in **Figure 6B**.

**Figure 6.**
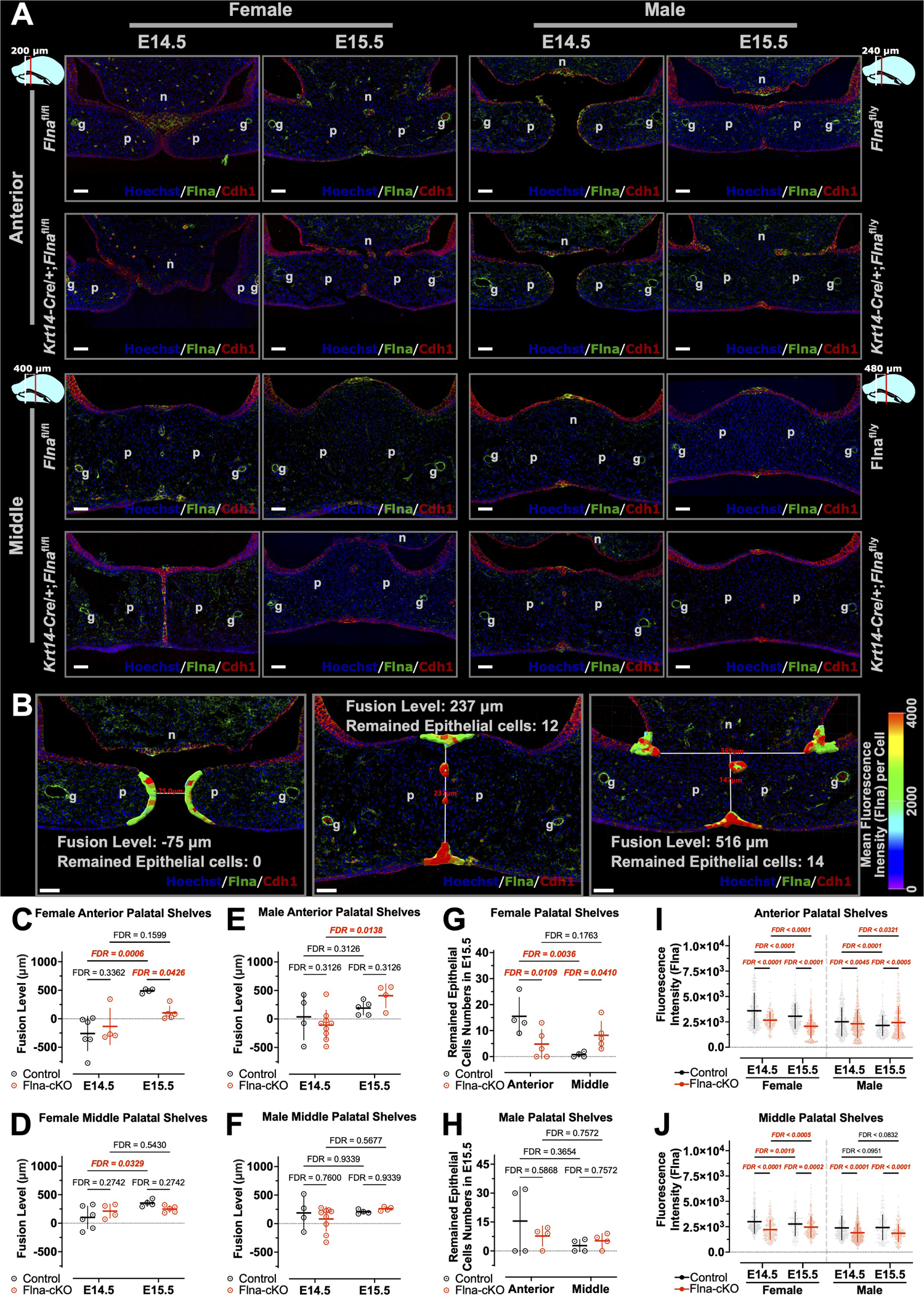
Epithelium-specific knockout of Flna delayed palatal fusion in female mice but not males. **A** Representative results of Flna-cKO in female/male anterior/middle palatal shelves at E14.5 and E15.5. **B**, **C, D, E, F** Illustration of the method to measure the “fusion level” and “remained epithelial cells numbers”. Flna-cKO induced changes of fusion level at female anterior (**C**), female middle (**D**), male anterior (**E**), and male middle (**F**) palatal shelves. **G, H** Flna-cKO induced changes in remaining epithelial cells numbers in male (**G**) and female (**H**) palatal shelves at E15.5. **I, G** Results of three-way analysis variance of fluorescence intensity of Flna from Min-Max normalized images in anterior (**I**) and middle (**J**) palatal shelves of both male and female from E14.5 to E15.5. **Data information: p**, palatal shelves; **n**, nasal septum; **g**, greater palatine artery; scale bar = 50 μm. The contrast, gamma, and brightness of all fluorescence images were uniformly adjusted for better presentation. All 2D images here are images after 3D projection. Quantitative results are presented by scatter plots showing individual data points along with mean ± standard deviation.

Control female mice showed a significant increase in fusion level of anterior palatal shelf from E14.5 to E15.5 (**Fig. 6C**). However, Flna-cKO mice showed no significant difference between these stages, leading to a notably lower fusion level at E15.5 (**Fig. 6C**). Although Flna-cKO was not a significant factor in the two-way analysis of variance, there was a significant interaction between developmental time and Flna-cKO in the fusion level changes of female anterior palatal shelves (**Table EV1**). This interaction suggests that effect of developmental time on fusion level was influenced by Flna-cKO, significantly delaying female anterior palatal shelf fusion from E14.5 to E15.5 (**Fig. 6C**). There were no significant differences in the fusion level of female middle palatal shelves between control and Flna-cKO mice, but Flna-cKO inhibited the fusion level in control mice (**Fig. 6D**). In control male mice, there were no significant increases in fusion levels of anterior and middle palatal shelves from E14.5 to E15.5 (**Figs. 6E** and **F**). Interestingly, male Flna-cKO mice showed a significant increase in anterior palatal shelf fusion (**Fig. 6E**).

The number of cells in the remaining medial epithelial islands of female control mice was significantly higher in the anterior than the middle palatal shelves at E15.5 (**Fig. 6G**). Flna-cKO eliminated this difference, resulting in fewer and more remaining MEE cells in the anterior and middle palatal shelves at E15.5, respectively (**Fig. 6G**). Despite insignificance of Flna-cKO in the two-way analysis of variance, there was a significant interaction effect between palatal shelf position and Flna-cKO on cell number changes in the remaining medial epithelial islands (**Table EV1**). This effect could suggest that Flna-cKO impacted the cell number difference between anterior and middle female palatal shelves (**Table EV1**). In both control and Flna-cKO male mice, there were no significant differences in remaining medial epithelial island cell numbers at E15.5 (**Fig. 6H**).

Notably, Flna-cKO significantly decreased Flna expression in all cases except male anterior palatal shelves at E15.5 (**Figs. 6I** and **J**). Consistent with the increased *Flna* expression in Flna-cKO male anterior palatal shelves at E15.5 (**Fig. 6I**), their fusion level also slightly increased (**Fig. 6E**).

### Parental habitual physical activity was causally associated with lower risk of cleft lip/palate

DNA sequence heterogeneity has been shown to influence human exercise behavior, with significant differences in the SNP profile between individuals with high and low exercise activity levels (Bouchard *et al*, 2011). The reported heritability of habitual physical activity (HPA) suggests that the embryonic SNP profile might also reflect parental exercise behavior (Hoed *et al*, 2013; Gielen *et al*, 2014; Klimentidis *et al*, 2018). To test whether HPA could causally affect the risk of cleft lip/palate (CLP), we conducted a two-sample Mendelian randomization (MR) analysis using GWAS summary statistics from the MRC IEU OpenGWAS database (Hemani *et al*, 2018).

The SNP characteristics related to HPA and CLP are depicted in **Figure 7A**, demonstrating that the SNPs were not weak instrumental variables (F = 18.11584) and were not directly associated with CLP, as no significant horizontal pleiotropy was detected (*P*-value = 0.42969), according to the intercept of the MR Egger regression (**Fig. 7A**). The causal effects of each SNP on CLP, showed in **Figures 7B** and **7C**, displays a symmetric pattern around the inverse-variance weighting (IVW) estimate in a funnel plot (**Fig. 7B**), suggesting that pleiotropy did not bias the causal effect.

**Figure 7.**
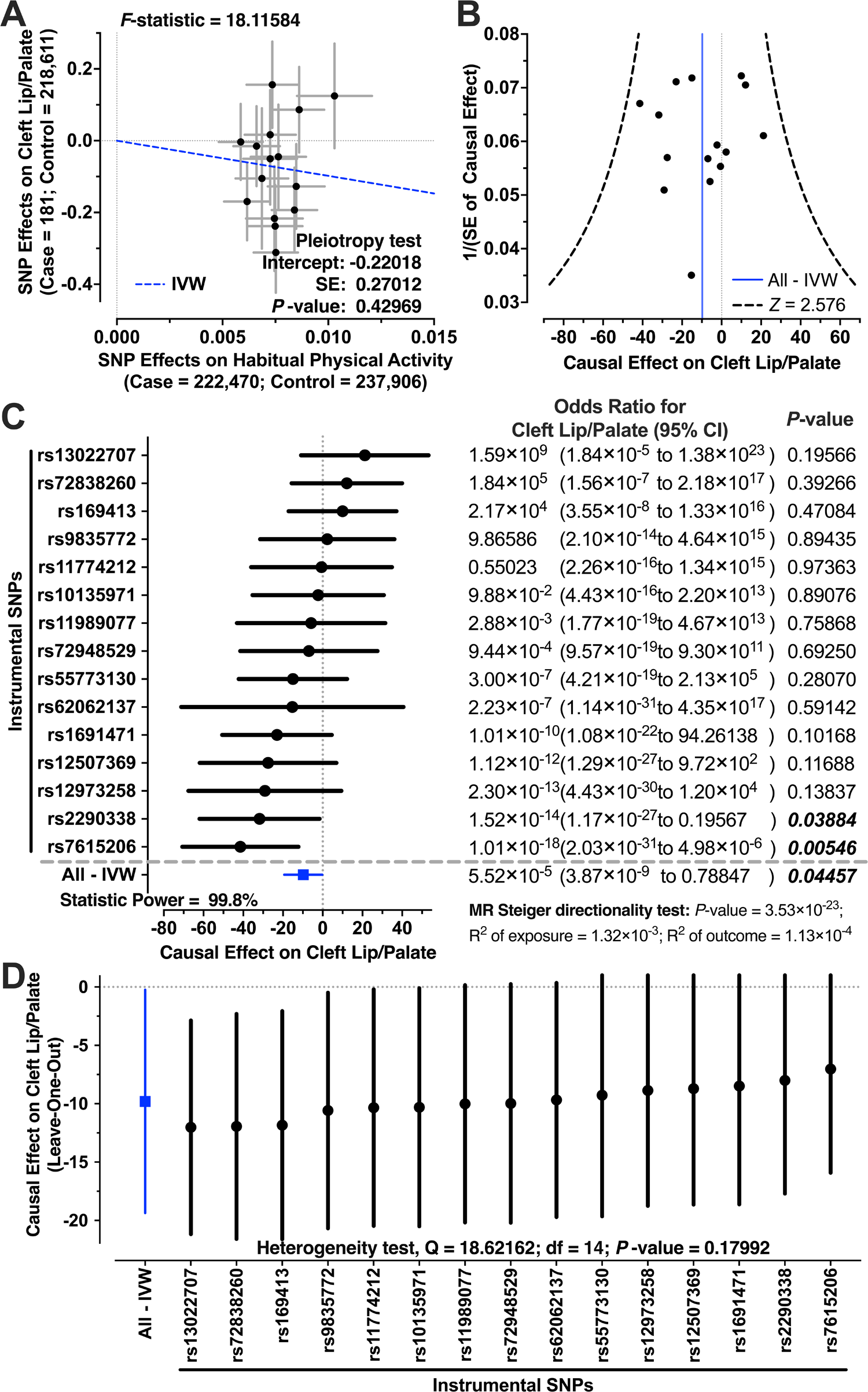
Parental habitual physical activity was causally associated with lower risk of cleft lip/palate. **A** Relationship between the effect size estimates on CLP (x-axis) and the effect size estimates on HPA (y-axis) for all SNPs that served as instrumental variables for CLP in the European population with the results of *F*-statistic and pleiotropy test using MR-egger intercept. Here, a total of 16 HPA instrumental variables were employed. The 95% confidence intervals for the estimated SNP effect sizes on CLP are shown as vertical black lines, while the 95% confidence intervals for the estimated SNP effect sizes on HPA are shown as horizontal black lines. **B** Funnel plot displays individual causal effect estimates for HPA on CLP in the European population. The dots represent the estimated causal effect for each IV. The vertical dotted red line represents the estimated causal effect obtained using all IVs from IVW estimate. **C** Forest plot for individual causal effect estimate with the results of MR Steiger directionality test and statistical power. **D** Leave-one-out plot to visualize causal effect of HPA on CLP risk when leaving one SNP out with the results of Cochran’s IVW Q test. **Data information: MR**, Mendelian randomization; **CLP**, cleft lip/palate; **HPA**, habitual physical activity; **SNP**, single-nucleotide polymorphism; **IVs**, instrumental variables; **IVW**, inverse-variance weighting. The code used for this MR analysis can be found in **Mendeley Data** (see **Reagents and Tools Table**).

From the IVW estimation, we observed that parental HPA was causally associated with the lower risk of CLP in their offspring, with an odds ratio (OR) of 5.52 × 10^-5^ (**Fig. 7C**). The 95% confidence interval ranged from 3.87 × 10^-9^ to 7.88 × 10^-1^, with a *P* -value of 0.04457 and a statistical power of 99.8% (**Fig. 7C**). The MR Steiger directionality test further supported the causality of parental HPA in reducing CLP risk (*P* -value = 3.53 × 10^-23^). As shown in **Figure 7D**, iterative MR analysis on the remaining SNPs after excluding each SNP in turn consistently supported the IVW estimation. Additionally, Cochran’s IVW Q test indicated no significant heterogeneity influence in this MR analysis (*P* -value = 0.17992).

## Discussion

In this study, we showed that parental exercise might reduce the risk of cleft palate in offspring, bridging clinical evidence with molecular biology mechanisms through MR analysis of large-scale GWAS datasets and genetic mouse models. Our findings demonstrate the crucial role of Flna in the TGF-β-regulated Smad-dependent and non-Smad-dependent pathways, such as RhoA during palatal fusion. Flna enhances TGF-β-induced β-catenin expression and responds to mechanical stimuli (**Figs. 5B** and **4D**), indicating its significance as a mediator in sustaining Wnt/TGF-β/Smad signaling activation in MEE cells during palatal fusion. Notably, only female Flna-cKO mice exhibited delayed palatal fusion in the anterior palatal shelves (**Figs. 6C-H**). Considering the established role of Wnt/TGF-β/Smad signaling in cleft palate (Marazita, 2012; Nakajima *et al*, 2018; Lan *et al*, 2015; Weng *et al*, 2018) and the higher prevalence of cleft palate in females (Pool *et al*, 2021), the X chromosome-linked gene, *Flna*, may partially explain this gender difference.

Global *Flna* knockout (Flna^Dilp2/y^ and Flna^Dilp2/Dilp2^) causes cleft palate by impeding palatal shelf elevation and results in prenatal lethality (Hart *et al*, 2006). To study role of *Flna* in palatal fusion, we used siFlna electroporation in *ex vivo* cultured palatal shelves at E14.0, after elevation of palatal shelves (Tarr *et al*, 2018). Although these shelves were of mixed genders from ICR mice, known for a lower cleft palate rate in TGF-β knockout (Sugiyama & Takigawa, 2020), the global knockdown of Flna still delayed palatal development more than Flna-cKO mice (**Figs. 4** and **6**). Our Flna-cKO mice, derived from C57BL/6J mice that have been shown the highest rate of complete cleft palate in TGF-β knockout (Sugiyama & Takigawa, 2020), showed no visible cleft palate in adult males, suggesting an unknown compensatory mechanism in MEE cells dependent on Flna in palatal mesenchymal cells.

Contrary to the original report (Feng *et al*, 2006), our *KRT14-Cre/+;Flna^fl/fl^*mice did not eliminate Flna proteins in MEE cells. Because these two mouse strains, *Flna*^tm1.1Caw^/J and Tg(*KRT14-cre*)1Amc/J, are widely used, and our previous work successfully deleted *Runx1* in MEE cells by using this Tg(*KRT14-cre*)1Amc/J strain (Sarper *et al*, 2018), this persistence in our Flna-cKO mice (**Fig. 6**) might be due to exogenous sources, like exosomes from neighboring mesenchymal cells or macrophages(Martínez-Álvarez *et al*, 2000), known for Flna-rich exosomes and proximity to MEE cells (Hassani & Olivier, 2013; Lee *et al*, 2009). Exosomes, containing various cleft palate-related substances (Chen *et al*, 2021), may play a role in cleft palate (Chen *et al*, 2022), though this is poorly reported.

A significant correlation between FLNA*/Flna* and β*-*catenin/*Ctnnb1* was observed in both HaCaT cells and EMT cells *in vivo* from scRNA-seq (**Figs. 5C** and **3E**). *Flna*-induced β-catenin/Smad2 changes in HaCaT cells and *ex vivo* cultured embryonic palatal shelves (**Figs. 4D** and **5**) suggest a general presence of Flna-regulated Wnt/Smad signaling in epithelial cells. However, the discrepancy of Flna-induced Smad2 changes between HaCaT cells and *ex vivo* cultured palatal shelves in our results suggested that this Flna-regulated Wnt/Smad signaling was subject to the regulation of local micro-environment, such as the epithelial-mesenchymal crosstalk (Lee *et al*, 2020).

Mendelian randomization (MR), an advanced statistical method, is employed to establish a causal relationship between an exposure of interest (e.g., HPA) and an outcome (e.g., the risk of cleft palate). This approach uses SNPs as IVs for the exposure. The underlying principle is that if HPA is causally linked to cleft palate, SNPs associated with HPA should also correlate with the risk of cleft palate through the path of HPA. Even if the selected SNPs are not directly causal for HPA but only show a high prevalence among individuals with regular HPA, MR can still elucidate the causal connection between HPA and cleft palate. The inheritability of SNPs suggests that the presence of HPA-related SNPs in offspring is likely inherited from parents who possess these SNPs, which are highly prevalent in populations engaging in regular HPA. In this study, our MR analysis suggested that European parents who exercise regularly can reduce the risk of cleft palate in their offspring by at least 21% (**Fig. 7C**). Consistent with our findings, Morton *et al*. (2013) reported that women with physical disabilities are at a higher risk of low birthweight and preterm birth (Morton *et al*, 2013), both of which are recognized risk factors for cleft palate (Wyszynski *et al*, 2004). The HPA-related SNP, rs7615206, exhibited the strongest causal effect on reducing CLP risk (*P* -value = 0.00546) and is located in MST1R gene. MST1R encodes the macrophage stimulating receptor, a cell surface receptor tyrosine kinase that transmits signals from the extracellular matrix into the cytoplasm through binding with its ligand, macrophage stimulating 1 (MST1). The MST1-regulated transduction process has been reported to be important in the mechanical regulation of bone remodeling (Wang *et al*, 2022). Notably, the activation and translocation of MST1 is regulated by the FLNA mechanobinding partner, SAV1 (Zhang *et al*, 2023).

Furthermore, our results showed that the expression of *Flna* in MEE cells was extremely sensitive to mechanical stimuli (**Figs. 2J-K**). The normal human walking may generate mechanical energy changes of 0.2 J/kg in each gait cycle (Tesio & Rota, 2019). The mass of the flask with 10 ml culture medium and 5 dissected *ex vivo* cultured palatal shelves was about 25 g in this study. In rough estimation, the rolling cultured flask received 4.5 × 10^-5^ J/kg, and the shaking cultured flask received 8.0 × 10^-5^ J/kg. A significantly increased *Flna* gene expression level in the MEE cells of palatal shelves was observed even if the total energy difference was only as slight as 3.5 × 10^-5^ J/kg (**Figs. 2J-K**). Since the *ex vivo* cultured palatal shelves were floating in the cultured medium, the actual energy delivered to each palatal shelf was even much smaller than this rough estimation. Thus, our study revealed that even slight mechanical stimuli can induce *Flna* expression in MEE cells, suggesting maternal exercise might benefit embryonic development. This aligns with recent findings on maternal exercise enhancing embryonic myogenesis (Gao *et al*, 2022).

This study suggested a potential mechanism linking mechanical stimuli to palatal development through a mechanosensor, Filamin A, and raised many unsolved problems related to the role of exosomes, epithelial-mesenchymal crosstalk, and exercise in palatal development. Therefore, given that exercise is one of the easiest accessible and non-invasive methods to improve individual health, further studies about relationship between mechanical stimuli and palatal development may be finally beneficial to make better maternity care and reduce the cleft palate risk in the future.

## Materials and Methods

### Re-analysis of publicly available bulk RNA-seq data along with literature mining

Original sequencing data from GSE45568 and GSE185279 were downloaded and quality trimmed (Liu *et al*, 2020; Ozturk *et al*, 2013; Sun *et al*, 2022). The reads were then mapped to the mouse genome (GRCm39, Ensembl release 106) with STAR aligner. The *de novo* transcripts assembly was performed using StringTie and was annotated using SQANTI3 software. All samples were then re-quantified at the transcript level using Salmon software with the *de novo* assembled transcriptome reference, and additional quality filtering was performed using isoformSwitchAnalyzeR based on the mapping results from Salmon. The significant differentially expressed genes (DEGs) were detected with DESeq2.

A literature mining (Cai *et al*, 2021) of the PubMed database was performed to extract the genes that highly correlated with the PubMed query results. A variable *R* was imported to evaluate the correlation between the input genes list and a PubMed query by calculating the ratio of (PubMed query and input gene-related articles)/(input gene-related articles). *P*-value was further applied to statistically analyze the chance probability of co-occurrences of each input gene and PubMed query in at least *K* papers, as following:

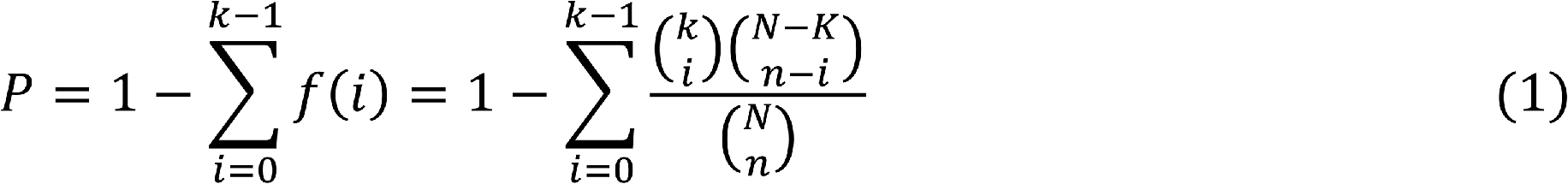

where *N* is the total number of articles in PubMed, *K* stands for the number of literatures related to input PubMed query term, n and k represent the amounts of articles for a specific gene and its corresponding articles on results from PubMed query, respectively. Subsequently, the nominal p-value was corrected as false discovery rate (FDR) using the Benjamini-Hochberg approach. Genes with *R* values greater than 10*K*/*N* and FDR or *P*-value (if *K* < 100) lower than 0.05 were considered as significance.

### Re-analysis of publicly available single-cell RNA-seq data

Original sequencing data from GSE155928 were downloaded (Han *et al*, 2021). The reads were aligned to the mouse genome (GRCm39, Ensemble release 106) and were counted by Cell Ranger software. Doublets were detected by using Solo software. Quality control, normalization, dimensionality reduction, clustering-based cell annotation, DEGs analysis, and visualization was performed using the Seurat R package. The RNA velocity analysis was performed using the scVelo Python module.

### HaCaT cells culture

HaCaT cells, an immortalized keratinocyte line from male human (RRID: CVCL_0038; AddexBio, San Diego, CA, USA), were cultured in 5% CO_2_ at 37 °C in regular DMEM containing 10% heat-inactivated fetal bovine serum, 100 U/mL penicillin, and 100 mg/ml streptomycin. Cells were seeded at 2.6 × 10^4^ cells/cm^2^ density and cultured with DMEM in 6-well culture palates. For experiments, the medium was supplemented with TGF-β3 (10 ng/mL; Proteintech, Rosemont, IL, USA) and/or 50 μM RhoA inhibitor (Y27632; MedChemExpress, Monmouth Junction, NJ, USA) and/or 75 pM siRNA or 0.1% DMSO for 48 hours.

### Palatal organ culture

The palatal shelves of ICR mice were cultured in suspension with roller or shaker for 24 or 48 hours, without gender determination. For dissection, E13.5 or E14.5 embryonic maxillary regions with palatal shelves were carefully separated using syringe needles (**Palates_Dissection.mp4** at **Mendeley Data**, see **Reagents and Tools Table**). Brain- and eye-forming tissues were removed, and three to five maxillary regions, including palatal shelves, were placed in 25 cm² rectangular flasks with vented caps containing 10 mL BGJb medium. These flasks were then set on tube rotators or shakers inside a sealed cabinet, which was flushed with 95% O_2_ and 5% CO_2_ to maintain oxygen levels above 80% during the culture.

For experiments involving chemical inhibitors, the medium was supplemented with either 20 μM TGF-β inhibitor (SB431542; MedChemExpress, Monmouth Junction, NJ, USA), 50 μM RhoA inhibitor (Y27632; MedChemExpress, Monmouth Junction, NJ, USA), or 0.1% DMSO for 24 hours.

In mechanical stimuli experiment, the tube rotator with15 rpm created a height change of 12 cm per rotation. The shaker operated at 120 Hz with a 4 cm amplitude. Considering the weight of culture flask (∼25 g), the energy of the system was estimated at approximately 4.5 × 10^-5^ J for the rolling group and 8.0 × 10^-5^ J for the shaking group, calculated using the formula of 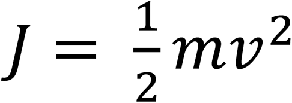.

### Mouse breeding

Mice with the targeted *Flna* gene, in which exons 3–7 of the Flna gene are located within loxP sites (*Flna^fl/fl^*), were obtained from the Jackson Laboratory (Strain #:010907; RRID: IMSR_JAX:010907). *KRT14*-*Cre* transgenic mice expressing Cre under the control of a human keratin 14 promoter/enhancer sequence were obtained from the Jackson Laboratory (Strain #:004782; RRID: IMSR_JAX:004782). Homozygous epithelium-specific Flna conditional knockout (Flna-cKO) mice in male (*KRT14-Cre/+;Flna^fl/y^*) generated from crosses of both mouse lines were backcrossed with *Flna^fl/fl^* mice, and the progeny were analyzed. Data obtained from homozygous floxed Flna littermates lacking *KRT14-Cre* were pooled and used as controls.

### Small interference RNA experiments

To knock down Flna in HaCaT cells and *ex vivo* cultured palatal organs using small interference RNA (siRNA), a stealth RNAi negative control medium GC duplex was purchased from Invitrogen (Carlsbad, CA, USA). Predesigned siRNAs targeting human FLNA (#2316-1, #2316-2, and #2316-3) and mouse Flna (#192176-1, #192176-2, and #192176-3) were purchased from Bioneer (AccuTarget™ predesigned siRNA; Daejeon, Korea). HaCaT cell transfection with a 75 pM siRNA mix (#2316-1, #2316-2, and #2316-3) was conducted using Lipofectamine RNAiMAX transfection reagent (Invitrogen, Carlsbad, CA, USA), following the manufacturer’s guidelines.

For *ex vivo* cultured palatal organ transfection, the dissected maxilla was held with an anode tweezertrode, and a cathode needle was positioned between two palatal shelves in the organ’s center. OPT-MEM containing a 75 pM siRNA mix (#2316-1, #2316-2, and #2316-3) was applied around the cathode needle between the palatal shelves (**Palates_Electroporation.mp4** at **Mendeley Data**, see **Reagents and Tools Table**). Electroporation was performed using a Super Electroporator NEPA21 Type II (Nepa Gene, Chiba, Japan), applying 80 V for poring (30 ms at 50 ms intervals, 3 pulses) and 70 V for transfer (50 ms at 50 ms intervals, 3 pulses).

### Frozen sections and immunofluorescence staining

The dissected embryonic heads from *ex vivo* cultured or *in vivo* samples were fixed with 4% paraformaldehyde in phosphate-buffered saline (PFA/PBS). The tissues were then sequentially immersed in sterile PBS solutions with increasing sucrose concentrations (10%, 20%, and 30%) until they sank. Subsequently, they were embedded in Tissue-Tek optimum-cutting-temperature (OCT) compound (Sakura Finetek, Tokyo, Japan). Sections of 6 μm thickness were cut from these OCT-embedded tissues, adhered to Kawamoto film (Kawamoto & Kawamoto, 2014), freeze-dried, and stored at −80°C. For reference, the anterior and middle palatal shelves were defined based on specific distances from the gap between primary and secondary palates at E14.5 and E15.5 stages (**E14_Palates_HE.pdf** at **Mendeley Data**, see **Reagents and Tools Table**).

For immunofluorescence staining, films stored at −80°C were thawed in PBS containing 0.1% Tween-20 (PBST) at room temperature. Tissues were permeabilized with 0.3% Triton in PBS for 3 minutes, re-fixed with 4% PFA for 15 minutes, and blocked with Blocking One Histo for 10 minutes at room temperature. They were then incubated overnight at 4 °C or two hours at room temperature with primary antibodies. Following three washes with PBST, films were incubated for 1 hour at room temperature with secondary antibodies and 1 μg/mL of DAPI or Hoechst 33342. After three additional PBST washes, the films were mounted and examined using a confocal laser scanning microscope.

### Quantitative analysis of immunofluorescence images

To analyze the epithelial triangle, 3D images were projected into 2D using “3D projection” function in Fiji/ImageJ software. Cells were manually outlined for morphology and fluorescence intensity analysis. Cell morphology was quantified using circularity, calculated as 4π×(Cell Area/perimeter^2^), and solidity, calculated as Cell Area/Convex Area.

For epithelial layer morphology, Cdh1-positive cells were identified as epithelial cells. The epithelium in the TGF-β inhibitor group was aligned to the control group by comparing epithelium thickness curves using Pearson’s correlation. The epithelium in the RhoA inhibitor group was aligned to the control group by the first discontinuation point of the epithelium layer. The epithelium was divided into upper (77–185 µm from the upper end), middle (epithelium disappeared in the control group), and lower (185 µm from the lower end) for cellular morphology and fluorescence intensity analysis. Upper and lower epithelial cells were identified and segmented using Imaris software.

For quantitative evaluation of palatal fusion, fusion level was defined as the distance between epithelial triangles if palatal shelves were conjunct (positive value) or between shelves if not conjunct (negative value). If shelves were also conjunct with the nasal septum, the fusion level was defined as the length of a line linking two epithelial triangles of the nasal septum to the palatal shelves and the distance from the palatal shelf triangle to this line. The remaining epithelial cell numbers were defined as the count of E-cadherin-positive cells between epithelial triangles if shelves were conjunct, or zero if not.

For fluorescence intensity comparison across samples, min-max normalization was performed [Intensity_Normalized_=(Intensity–min(Intensity))/(max(Intensity)–min(Intensity))]. This normalization preserved relative fluorescence intensity strength of Flna, β-catenin, Smad2, E-cadherin, and Hoechst 33342 between siControl and siFlna, making relative intensities of these genes comparable across samples. For 3D confocal images of Flna-cKO samples, minimal intensity was background fluorescent noise, and maximal Flna intensity was from the greater palatine artery (**01.All_analysis_results_of_80_images.pdf** in **Mendeley Data**, see **Reagents and Tools Table**). This approach preserves relative Flna intensity in MEE cells compared to background and the greater palatine artery, ensuring comparability across images.

All image processing and analysis were conducted using Fiji/ImageJ, Imaris software, and the R programming language. Original images, ImageJ macro scripts, and R scripts are available in **Mendeley Data** (see **Reagents and Tools Table**).

### Western blotting

HaCaT cells were washed with cold PBS and lysed, followed by sonication on ice. Protein concentrations were determined and adjusted to 0.29 mg/mL. Four micrograms of each protein sample were separated using polyacrylamide gel electrophoresis. The proteins were then transferred to polyvinylidene difluoride (PVDF) membranes. PVDF membranes were blocked in a solution of 5% nonfat skim milk in PBST for 1 hour, followed by incubation for 1 hour at room temperature with primary antibodies. After three 10-minute washes in PBST, the membranes were incubated with corresponding secondary antibodies for 1 hour at room temperature. After three additional washes, proteins were detected using the ChemiDoc™ XRS+ system (Bio-Rad, Berkeley, CA, USA) with 20X LumiGLO^®^ Reagent and 20X Peroxide (Cell Signaling, Danvers, MA, USA).

Densitometric analysis of the bands was performed using ImageLab software (version 6.0.1; Bio-Rad, Berkeley, CA, USA). The molecular weight (MW) of the target proteins was estimated by a standard curve, plotting the logarithm of MW against relative migration distance, generated using Precision™ Plus Protein Dual Xtra Standards.

### Mendelian randomization analysis

Genetic variants for habitual physical activity (HPA) and cleft lip/palate (CLP) were obtained from MRC IEU OpenGWAS database (Hemani *et al*, 2018) with ID of “ukb-b-8764” and “finn-b-Q17_CLEFT_LIP_CLEFT_PALATE”, respectively. The following selection criteria were used to choose the instrumental variables (IVs): (1) single nucleotide polymorphisms (SNPs) associated with each genus at the locus-wide significance threshold (*P* < 5.0×10^-8^) were selected as potential IVs; (2) 1000 Genomes project European samples data were used as the reference panel to calculate the linkage disequilibrium (LD) between the SNPs, and among those SNPs that had *R*^2^ < 0.001 (clumping window size=10,000 kb), only the SNPs with the lowest *P*-values were retained.

Two-sample MR analysis was performed based on the “TwoSampleMR” R Package, and inverse variance weighting (IVW) was used to assess the causal association between HPA and the risk of CLP. The strength of IVs was assessed by calculating the *F* -statistic using the formula 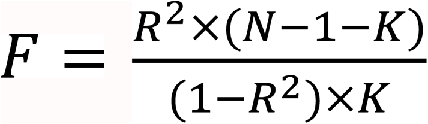, where *R*^2^ represents the proportion of variance in the exposure explained by the genetic variants, *N* represents sample size, and *K* represents the number of instruments. If the corresponding F-statistic was >10, it was considered that there was no significant weak instrumental bias(Instrumental Variables Regression with Weak Instruments, 2024). The power of the MR estimates was calculated using the online calculator tool provided by Stephen Burgess (Burgess, 2014). The causal direction between the hypothesized exposure and outcomes can be tested using the Steiger test (Hemani *et al*, 2017). Cochran’s IVW Q statistics were used to quantify the heterogeneity of IVs. In addition, to identify potential heterogeneous SNPs, the “leave-one-out” analysis was performed by omitting each instrumental SNP in turn. The intercept of MR-Egger regression was used to evaluate the directional pleiotropy of IVs (Burgess & Thompson, 2017). *Z*-score of casual effect sizes of IVs was used to identify the outliers.

### Statistical analysis

For re-analysis of both bulk RNA-seq and scRNA-seq, DEGs were defined as FDR < 0.05 & |log2Fold Change| > 1, and over-representation analysis was conducted using the Gene Ontology (GO) database by clusterProfiler 4.0.

The normality was tested using the Shapiro-Wilk test with α set at 0.01. For comparing multiple groups, we used a one-way analysis of variance (ANOVA) followed by a multiple comparisons test with false discovery rate (FDR) control for normally distributed data (i.e., results from Western Blotting). For non-normally distributed data (i.e., results from the inhibitor experiments), the Kruskal-Wallis test followed by a multiple comparisons test with FDR control was employed. The Mann-Whitney test was used to determine statistical differences in normalized fluorescence intensity of Flna, β-catenin, Smad2, E-cadherin, Hoechst 33342, and cellular sphericity between siControl and siFlna in *ex vivo* cultured palatal shelves. The paired *t*-test was utilized for comparing mean Upper-Epi thickness and epithelial disappearing length between siControl and siFlna in these shelves.

Spearman’s coefficient was calculated for results from inhibitor experiments and analysis of the epithelial triangle, while Pearson’s coefficient was used for re-analysis of scRNA-seq and Western Blotting, to determine correlation strengths. Given the robustness of two-way and three-way ANOVA against normality violations (Blanca *et al*, 2017), normality testing for these analyses was not performed.

All statistical analyses were conducted using GraphPad Prism software (version 8.0.0 for MacOS, GraphPad Software, Inc., San Diego, CA, USA) or the R programming language. Quantitative results are presented in scatter or violin plots showing individual data points, accompanied by mean ± standard deviation for normally distributed data or median with interquartile range for non-normally distributed data. Regression lines are shown with 95% confidence intervals, and linked lines of paired data are presented for the paired *t*-test.

### Ethical considerations and animals

Protocols, handling, and care of the mice conformed to the guidelines of the Okayama University Committee for Care and Use of Laboratory Animals and were approved (OKU-2020705 for *ex vivo* culture of palatal shelves, and OKU-2022840 for Flna-cKO mice) by the Animal Experiments Ethics Committee of Okayama University. All mouse employed in this study were maintained on a 12 h/12 h light/dark cycle at between 23–26 °C and housed in cages with access to food and *ad libitum* autoclaved water.

## Data availability

The representative confocal laser scanning images, all original results of the image analysis, the original images of Western Blotting, the details of Western Blotting analysis, the R code to reproduce all bioinformatics analysis in this study, and the ImageJ macro scripts for image processing have been deposited at Mendeley Data, and all original confocal laser scanning images of biological replicates in .czi format were deposited at Figshare. These data are publicly available as of the date of publication. Accession numbers are listed in the **Reagents and Tools Table**.

We made a website (https://djlm3g1x1t4ic.cloudfront.net) to host the re-analysis results of bulk RNA-seq data from GSE45568 and GSE185279, as well as scRNA-seq data from GSE155928. This comprehensive re-analysis includes results from alternative splicing analysis results, circle RNA detection results, co-expression analysis, functional enrichment analysis results, DEG analysis results, and more (https://djlm3g1x1t4ic.cloudfront.net/BulkRNAseq/00.Analysis_Report.html). The re-analyzed scRNA-seq of GSE155928 are presented through an interactive web application that allows the user to check the UMAP plots, check the gene expression, make dot plot, make heatmap, do clustering analysis, do DEGs analysis, and more (https://dwll26k42dcbb.cloudfront.net/GSE155928/).

## Acknowledgments

This study was supported by the Japan Society for the Promotion of Science, Japan, through Grants-in-Aid for Scientific Research to Ziyi Wang (22F22114), Satori Hayano (22K10245), Mitsuaki Ono (22H03280), Toshitaka Oohashi (22KF0265 and 19K09649), and Hiroshi Kamioka (22K19624). We express our gratitude to Dr. Tomoko Yonezawa and his colleagues at the Department of Molecular Biology and Biochemistry, Okayama University. Special thanks to Mr. Tetsushi Iwasa from the Central Research Laboratories, Okayama University Medical School, for his extensive assistance with the ZEISS LSM 780 confocal laser scanning microscopy system.

## Author Contributions

**Ziyi Wang, Satoru Hayano**: Conceptualization, Methodology, Resources, Funding acquisition, Investigation, Formal analysis, Data curation, Visualization, Writing - Original draft, Writing - Reviewing and Editing. **Ziyi Wang**: Software, Bioinformatics. **Jeremie Oliver Piña**, **Rena N. D’Souza**: Writing - Original draft, Writing - Reviewing and Editing. **Mitsuaki Ono**, **Toshitaka Oohashi**, **Hiroshi Kamioka**: Funding acquisition, Methodology. **Yao Weng**, **Xindi Mu**, **Takashi Yamashiro**: Methodology, Investigation. **Yao Weng**, **Xindi Mu**, **Mitsuaki Ono**, **Toshitaka Oohashi**, **Hiroshi Kamioka**: Investigation, Formal analysis. **Ziyi Wang**, **Satoru Hayano**, **Rena N. D’Souza**, **Toshitaka Oohashi**, **Hiroshi Kamioka**: Supervision.

## Expanded View Tables

**Table EV1.** The results of two-way and three-way analysis of variance.

